# Adaptive Tuning Curve Widths Improve Sample Efficient Learning

**DOI:** 10.1101/775163

**Authors:** Florian Meier, Raphaël Dang-Nhu, Angelika Steger

## Abstract

Natural brains perform miraculously well in learning new tasks from a small number of samples, whereas sample efficient learning is still a major open problem in the field of machine learning. Here, we raise the question, how the neural coding scheme affects sample efficiency, and make first progress on this question by proposing and analyzing a learning algorithm that uses a simple reinforce-type plasticity mechanism and does not require any gradients to learn low dimensional mappings. It harnesses three bio-plausible mechanisms, namely, population codes with bell shaped tuning curves, continous attractor mechanisms and probabilistic synapses, to achieve sample efficient learning. We show both theoretically and by simulations that population codes with broadly tuned neurons lead to high sample efficiency, whereas codes with sharply tuned neurons account for high final precision. Moreover, a dynamic adaptation of the tuning width during learning gives rise to both, high sample efficiency and high final precision. We prove a sample efficiency guarantee for our algorithm that lies within a logarithmic factor from the information theoretical optimum. Our simulations show that for low dimensional mappings, our learning algorithm achieves comparable sample efficiency to multi-layer perceptrons trained by gradient descent, although it does not use any gradients. Furthermore, it achieves competitive sample efficiency in low dimensional reinforcement learning tasks. From a machine learning perspective, these findings may inspire novel approaches to improve sample efficiency. From a neuroscience perspective, these findings suggest sample efficiency as a yet unstudied functional role of adaptive tuning curve width.

## 1 Introduction

Humans operate in a rich and complex world and are extremely fast in learning new tasks and adapting to new environments. The level of generalization and speed of adaptation achieved by human brains remain unmatched by machine learning approaches, despite tremendous progress in the last years. How do real brains accomplish this outstanding skill of generalization and sample efficient learning, and what are the neural mechanisms that contribute to this ability of fast learning? Here, we investigate how neural coding supports sample efficient learning, by analyzing a learning algorithm that exploits three bio-plausible principles for sample efficient learning, namely, population codes of tuned neurons, continuous attractor mechanisms and probabilistic synapses.

From early on, neuroscience researchers characterized the first order response of single neurons by neural tuning curves [1]. The *neural tuning curve* is defined to be the neurons mean firing rate as a function of some stimuli parameter. It typically peaks for a preferred parameter value and decays gradually as this parameter moves away from the preferred value, such as in orientation columns in the visual cortex [2, 3], spatially tuned cells in auditory cortex [4], direction selective cells in motor cortex [5] and hippocampal place and head direction cells [6, 7]. In populations of tuned neurons, narrow (broad) tuning curves imply that a small (large) fraction of neurons is active for a given stimuli parameter. Here, we assume that such neural populations are geometrically ordered according to the neuron’s preferred parameter value. Then, the neural activity resembles a localized bump activation like experimentally observed in the compass system of the drosophila fly [8, 9] and theoretically studied in continuous attractor models [10, 11, 12, 13, 14]. In this coding scheme, which we call *bump coding scheme*, the center of a bump activation corresponds to the parameter value encoded by the bump activation (see Figure 1A), and the width of the bump is determined by the number of active neurons in the bump. Further, we assume that neural populations are equipped with a *continuous attractor mechanism* that ensures that only one bump is active at a time. Continuous attractor mechanisms are an established model of cortical working memory [14], and emerge from the wiring motive of local excitation and long range inhibition [9]. As another bio-plausible ingredient, we use *probabilistic synapses*. We assume a simple synaptic model consisting of a plastic synaptic probability *p* and a synaptic weight *w*, which we fix to 1 in order to concentrate on our main ideas. The synaptic probability corresponds to the pre-synaptic neuro-transmitter release probability and the weight *w* to the post-synaptic quantal amplitude [15]. The neuro-transmitter release probability of synapses in the brain is highly variable and typically between 0.1 and 0.9 [16].

**Figure 1:**
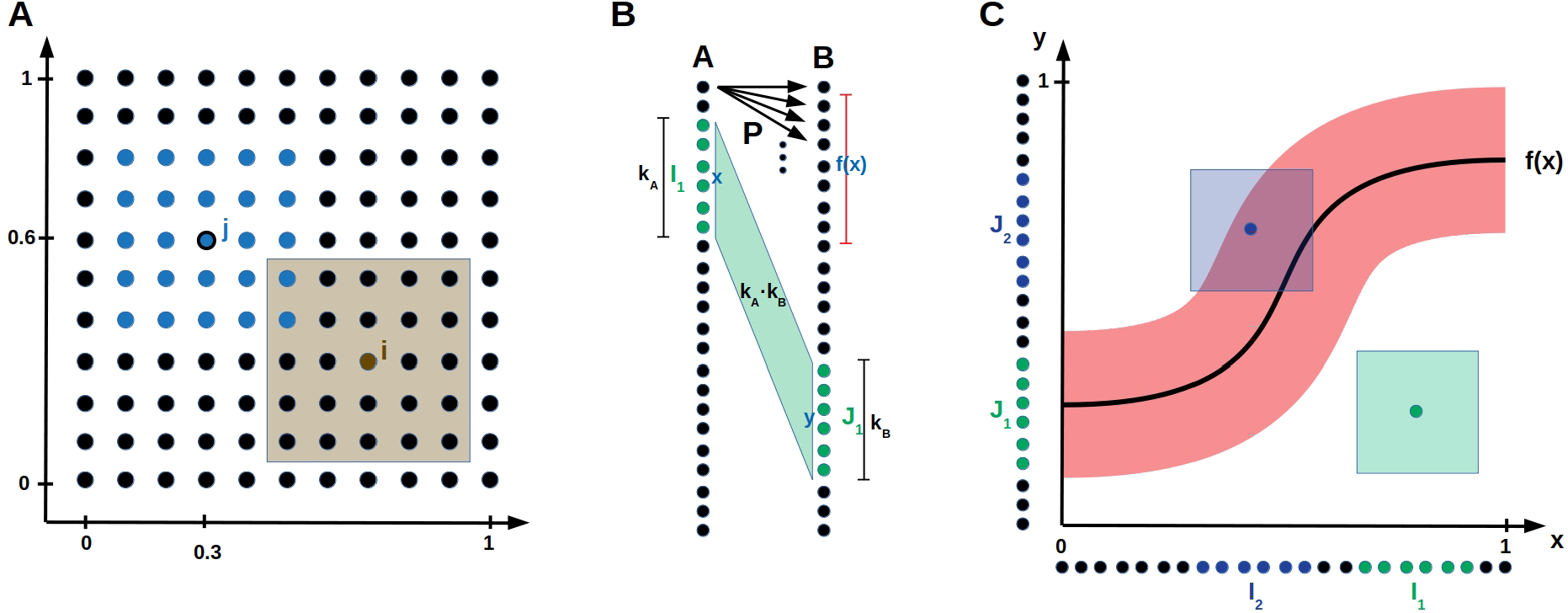
Bump coding scheme and model setup. (A) A neural population encoding a 2-dimensional parameter. The dots represent neurons. The blue neurons indicate a bump activation with center neuron *j* encoding the value (0.3, 0.6). The brown shaded area visualizes the parameter values for which neuron *i* is active. (B) Network setup. Two populations *A* and *B* of neurons both encoding a one-dimensional parameter are connected by probabilistic synapses *P*. The green input bump *I*_1_ with center *x* and width *k*_*A*_ activated the green output bump *J*_1_ with center *y* and width *k*_*B*_. The shaded area between the two bumps visualizes the *k*_*A*_ · *k*_*B*_ synapses that are updated according to the error feedback *L*(*x*, *y*). ((C) The neural populations *A* and *B* are aligned on the *x*- and *y*-axis, respectively, such that the synapses connecting population *A* to *B* are visualized by the (*x*, *y*)-space. The red area around the function *f*(*x*) indicates for each input *x* the *y* values that receive error feedback *L* smaller than some error threshold 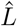. If an input output pair (*x*, *y*) is inside the red area (e.g. for the blue sample activation), then the theoretical bump algorithm increments the synaptic counters of the synapses between the bumps. Otherwise (e.g. for the green sample activation) it decrements the synaptic counters for the synapses between the bumps.

We explain now with a sample application, how these three bio-inspired principles are integrated into a reinforce-type [17], gradient-free learning mechanism. Assume that a robot arm with two joints should learn to reach given target positions *x* = (*x*_1_, *x*_2_) by applying the correct angles 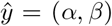 to the joints, such that they reach the given target position *x*. Consider two populations of neurons *A* and *B* connected by probabilistic synapses. Population *A* and *B* use the bump coding scheme to encode the target position *x* and the angles *y* that are applied to the two joints, respectively, see Figure 1B. The goal is to adapt the synaptic probabilities such that every target position *x* is mapped to the correct angles 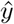. A bump activation encoding *x* in population *A* is propagated via probabilistic synapses to population *B*, where an abstract continuous attractor mechanism ensures that a single bump remains active in *B*. Its center *y* encodes the applied angles (*α*, *β*). Note that *y* usually varies from trial to trial since the synapses between population *A* and *B* are probabilistic. According to the final arm position, the network receives a scalar error feedback *L*(*x, y*), that depends on the input *x* and the output *y* generated by the network. Then, the synaptic probabilities between the bumps in population *A* and *B* are updated according to this error feedback. The larger the bump width *k* is, the more synapses are between the two bumps, whose probabilities are all updated according to the same error feedback. In this way, the network exploits the continuity of the task for sample efficient learning.

Related to this work, populations of tuned neurons have been used to learn sensory-motor transformations [18, 19, 20, 21, 22], extending and building on the investigation of radial basis networks conducted in the 1980s and 1990s [23, 24, 25, 26, 27, 28]. These studies use Hebbian plasticity mechanisms, that require simultaneous activation of inputs and target outputs, whereas our learning algorithms only rely on scalar error feedback. Furthermore, the investigation of the relation between sample efficiency and tuning curve width is novel.

The main contributions of this paper are summarized as follows. We introduce a reinforce-type learning algorithm that exploits the bump coding scheme, abstract continuous attractor mechanisms and probabilistic synapses for sample efficient learning of general low dimensional mappings. We show theoretically and by simulations that if the bump width is static during learning, then large bump width improves sample efficiency but harms the final precision, whereas small width impairs sample efficiency but improves final precision. Benefits of both are accomplished, if the bump width is dynamically decreased during the learning progress. Moreover, we show that the obtained sample efficiency is asymptotically optimal up to a log *n* factor in the limit of large population size *n*. For low dimensional mappings, the bump coding scheme achieves similar performance as a multi-layer perceptron trained by the backpropagation algorithm [29], and it outperforms a multi-layer perceptron trained by the reinforce algorithm [17]. It also achieves competitive performance on low dimensional reinforcement learning environments. Finally, we relate our findings to experimental observations of decreasing tuning curve width during learning and conclude that our findings propose sample efficiency as a functional role of the tuning curve width.

## 2 Results

Assume that a network consisting of populations *A* and *B* should learn a mapping 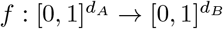, e.g. mapping target position 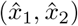 to joint angles (*α*, *β*) as illustrated in the robotic arm task of the introduction. We consider general mappings *f*, that are only restricted to be Lipschitz continuous^1^. This general framework, can be applied to many tasks including reinforcement learning as demonstrated in Section 2.2. Populations *A* and *B*, are connected by probabilistic synapses and encode input *x* and output *y*, respectively, using a bump coding scheme, see Figure 1 and Section 3.1 for a formal description. The goal is to learn the plastic synaptic probabilities, whereas synaptic weights are assumed to be fixed. The implicit goal of our learning algorithm is that a neuron *x* in population *A* keeps all synapses to the neurons *y* in *B* for which |*y* − *f*(*x*) is small and decreases the synaptic probabilities of all other synapses, see Figure 1C. This will ensure that a bump *x* in population *A* activates a bump with center *y* close to *f* (*x*) in population *B*.

We begin by stating our theoretical results in Section 2.1 before presenting the results obtained by simulations in Section 2.2. For the theoretical results, we use a simplified version of the algorithm used in the simulations, because it allows a rigorous mathematical analysis and it illustrates the conceptual ideas of the algorithm. Both algorithm use the same basic principles and behave qualitatively the same.

### 2.1 Theoretical Results

We consider the following learning mechanism with fixed bump width *k* involving synaptic counters that are initialized with 0. We refer to a neuron with preferred parameter value *x* as neuron *x* and to a bump with center *x* as bump *x*. For every sample, a random input bump *x* is activated. The probabilistic synapses propagate the activity in *A* to population *B*, where an abstract continuous attractor mechanism activates the bump *y* in *B* that received highest synaptic input. Then, the counters of the synapses between the two active bumps are decremented by 1 if the error feedback *L*(*x*, *y*) is larger than some error threshold 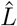 and incremented otherwise, see Figure 1C. After observing proportional 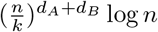 many samples, we prune all synapses with non-positive counters, that is, we set their synaptic probabilities to 0. For a formal description of the algorithm, we refer to Section 3.

We define the *error* of a learned network to be the expected error feedback 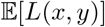 if the input is randomly chosen. If the mapping *f* is Lipschitz continous and the network obtains the euclidean distance *L*(*x*, *y*) = ‖*y* − *f*(*x*)‖_2_ as error feedback, the following theorems hold.

#### Theorem 1 (Static bump width *k*).

*The learning algorithm with static bump width k and euclidean error feedback learns a mapping* 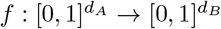 *with error smaller than* 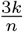 *after proportional to* 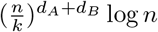 *many random samples, where n is the population size*.

The approach with static bump width *k* ensures that each neuron *x* maintains the synapses to a small continuous interval of output neurons around value *f*(*x*) and prunes away the other synapses. Thus, we can reapply the same learning mechanism, this time with smaller bump width *k* and smaller error threshold 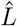 (for a formal description see Section 3). Repeating this procedure will cause an input bump *x* to be mapped to a random bump *y* from a shrinking and shrinking interval around *f*(*x*). This yields the following theorem.

#### Theorem 2 (Dynamic bump width *k*).

*The learning algorithm with dynamic bump width and euclidean error feedback learns the mapping* 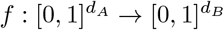 *with error smaller than ε after proportional to* 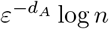 *many random samples, where n is the population size*.

We conclude that in order to reach error 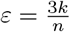 the dynamic bump algorithm requires proportional to 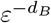 times less samples than the static bump algorithm with bump width *k*. Our proofs show that it is not necessary that above algorithms obtain the precise Euclidean distance as error feedback, but rather one bit of feedback suffices, if it indicates whether the Euclidean distance is larger than the error threshold 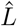 or not. For such an algorithm, a lower bound on the sample efficiency can be obtained by an entropy argument.

#### Theorem 3 (Lower Bound).

*For any algorithm that obtains only a single bit of feedback per sample, there are Lipschitz continuous mappings* 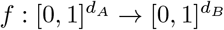, *such that the algorithm requires at least proportional to* 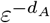 *many samples to learn f with error smaller than ε*.

Then, Theorem 5 and 6 imply that the learning mechanism with dynamic *k* accomplishes a sample efficiency that is asymptotically optimal up to a log *n* factor. For the proofs of these theorems, we refer to the supplementary material.

### 2.2 Empirical results

For the simulations, we use a slightly more sophisticated learning algorithm that follows the same underlying principles, but differs from the algorithm that we analyze theoretically in five aspects. Firstly, it is designed to handle more general error feedback functions. For example, in the robotic arm task of the introduction, the error feedback is not given by the euclidean distance between the output angles, but by the distance between the reached position and the target position. In turn, the magnitude of the error feedback can change for different inputs, and a single error threshold 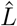 will not allow fast learning for all inputs. To resolve this issue, we assume that every input neuron *i* keeps track of a running mean 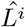 of the error feedbacks that were obtained when *i* was active. For the update of the outgoing synapses of neuron *i*, 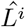 is compared to the error feedback *L*. Secondly, for every neuron *i* of the bump in population *A*, the synapses projecting on the bump in population *B* are pruned away immediately after observing a sample with 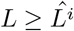. Thirdly, for the dynamic case, the bump width *k* is adapted continuously instead of repeatingly applying the static algorithm. Since the learning progress might vary for different input regimes, we allow *k* to depend on the input, and we set *k* for input *x* proportional to the number of outgoing synapses of neuron *x*. Note that this number is a reasonable measure of how well input *x* is already learned as the precision of the output depends on the magnitude of the interval of synapses connecting *x* to neurons around *f* (*x*). Fourthly, long-time inactive synapses are pruned away, i.e. they are pruned if the post-synaptic neuron has not been active for a couple of times when the pre-synaptic neuron was active. Finally, synapses of neuron *i* are consolidated (that is its probability is set to 1) if its number of synapses drops below a certain threshold value. We call this algorithm the *dynamic bump algorithm*. **W**e refer to Section 3 for a formal description of the algorithm and to Figure 2 for an illustration of the evolution of the synaptic probabilities.

**Figure 2:**
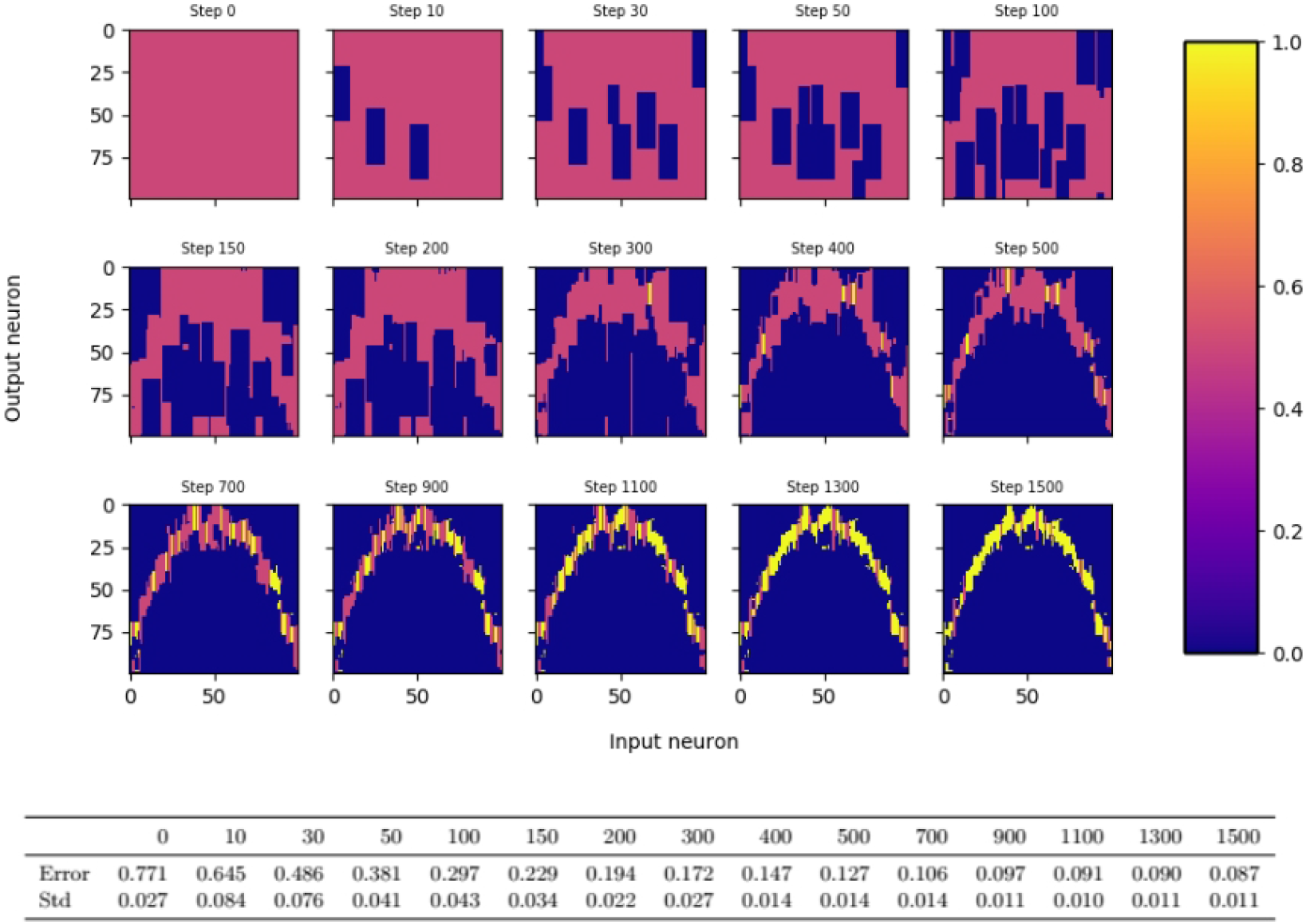
Evolution of synaptic probabilities when learning a one dimensional mapping *f*(*x*) = *x*^2^ − 3*x* + 1 with the dynamic bump algorithm. Each small color plot displays the synaptic probabilities, where input neuron is on the *x*-axis and output neuron on the *y*-axis. Blue areas visualize pruned synapses, yellow areas visualize consolidated synapses. Input and output population consist of 100 neurons each and the output bump is 3 times as large as the input bump.

The *static bump algorithm* works analogously to the dynamic bump algorithm, except that *k* is held constant during the whole algorithm. Figure 3 empirically confirms the trade-off between sample efficiency and final performance for static bump width *k*. Larger *k* leads to faster learning compared to smaller *k*, however reaches worse final error. The advantages of both large and small *k* can be exploited by adapting the bump width dynamically during the learning process, see Figure 3.

**Figure 3:**
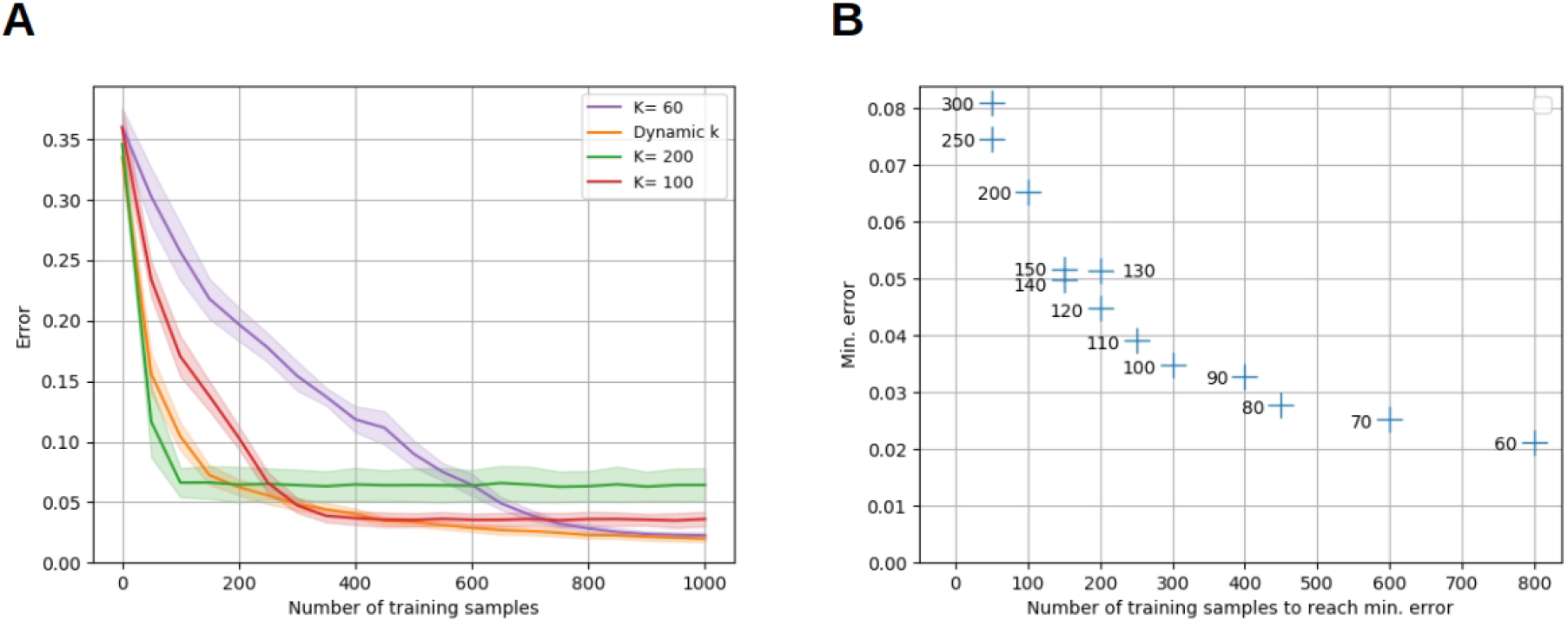
Sample efficiency versus final error trade-off for static bump width *k*. Recall that the error is defined to be the expected error feedback if the input is randomly chosen. Plot (A) shows the evolution of the error of the static bump algorithm for different *k*, when learning the one-dimensional identity mapping *f*(*x*) = *x* with euclidean error feedback. The dynamic bump algorithm, is labeled as ‘dynamic *k*’. Plot (B) shows the minimal error against the number of training samples required to reach this error for different *k*; to avoid taking into account the slow progress before final convergence, the number of samples required to achieve 1.5 times the final error is plotted. For both plots, populations *A* and *B* consist of 1000 neurons each, and the mean of 10 trials is plotted.

In order to put the sample efficiency of the bump coding scheme into context with other coding schemes, we compare it to the performance of a multi-layer perceptron (MLP), which encodes information with real valued units. Figure 4 compares performance of the dynamic bump algorithm, with a MLP trained by the backpropagation algorithm [29] and a reinforce algorithm as described in [17]. Note that the backpropagation algorithm requires full access to the first order derivative of the error with respect to the parameters and thus is a first-order optimization technique, whereas the reinforce and dynamic bump algorithm only require a scalar error feedback and thus are zeroth-order optimization techniques. Nevertheless the dynamic bump algorithm achieves similar performance as the MLP trained by backpropagation and outperforms the MLP trained by the reinforce algorithm, Figure 4. For the backpropagation and reinforce algorithm, we used a hyper-parameter search to determine the best parameters. **W**e note that this search yielded an untypically high learning rate and small batch size for the backpropagation algorithm. The learning rate is in the upper end of the recommended interval [10^−6^, 1] and much higher than the suggested default value of 0.01 [30]. This is necessary to achieve good performance after 1000 samples, see Figure 4.

**Figure 4:**
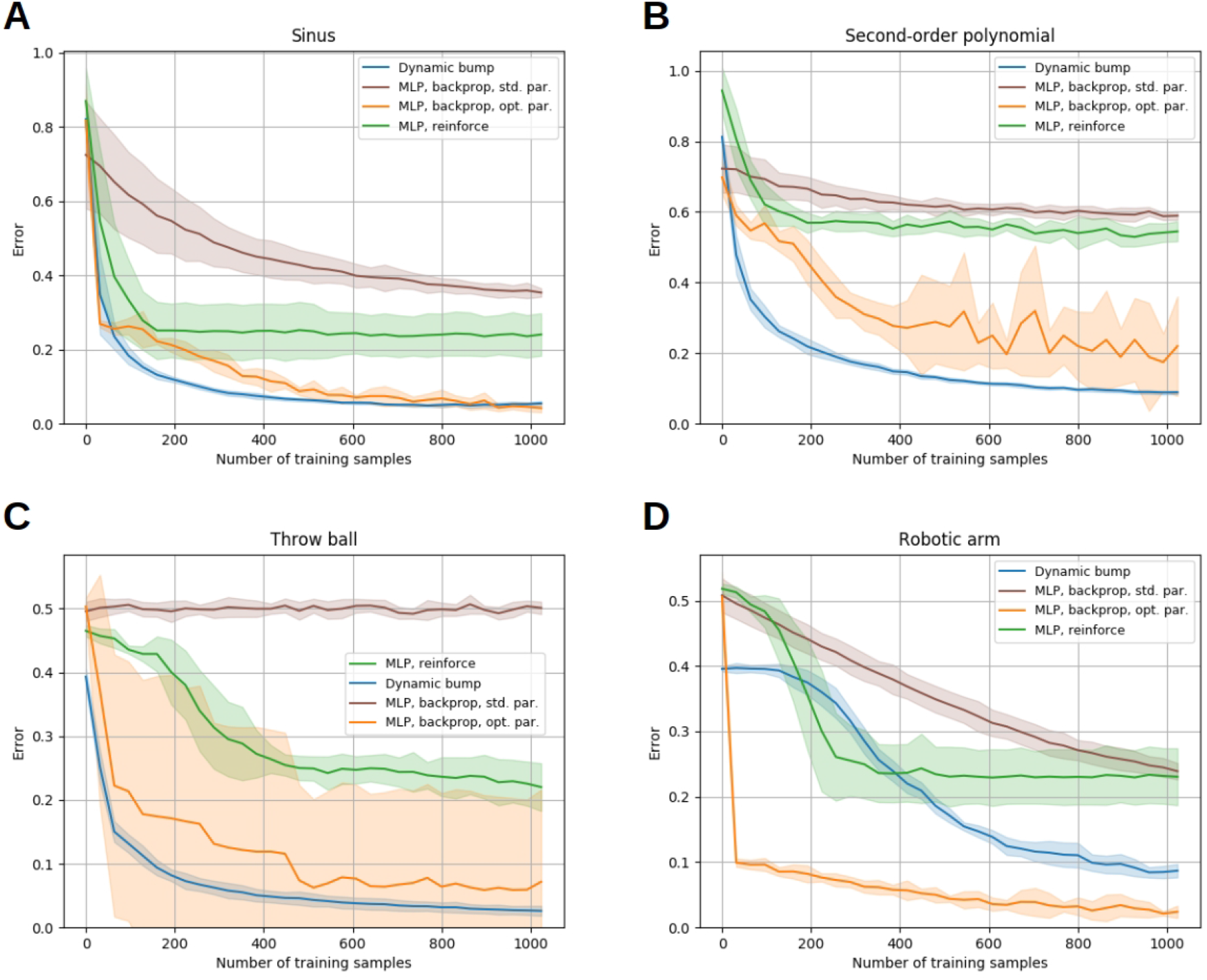
Comparison of the sample efficiency to a multi-layer perceptron. The learning progress of the dynamic bump algorithm, of a MLP trained with the backpropagation algorithm and of a MLP trained with the reinforce algorithm is shown for four tasks. The tasks are described in detail in Section 3.5. We plot the learning curve that achieves best performance after 1000 samples obtained by our hyper-parameter search. For each algorithm, we plot average and standard deviation of 10 runs.

A nice additional property of our algorithm is that the bump algorithm does not suffer from forgetting, if learning samples are ordered arbitrarily. We illustrate this in Figure 5 for the throw ball task, see Section 3.5. **W**hen training first only on samples with short distance targets and then only on samples with long distance targets, the performance on short distance targets is not hampered in the second phase, see Figure 5. Learning the solution for long distance targets does not interfere with the already learned solution for short distance targets because these solutions are stored in different synapses except for inputs close to the boundary of short and long distance inputs.

**Figure 5:**
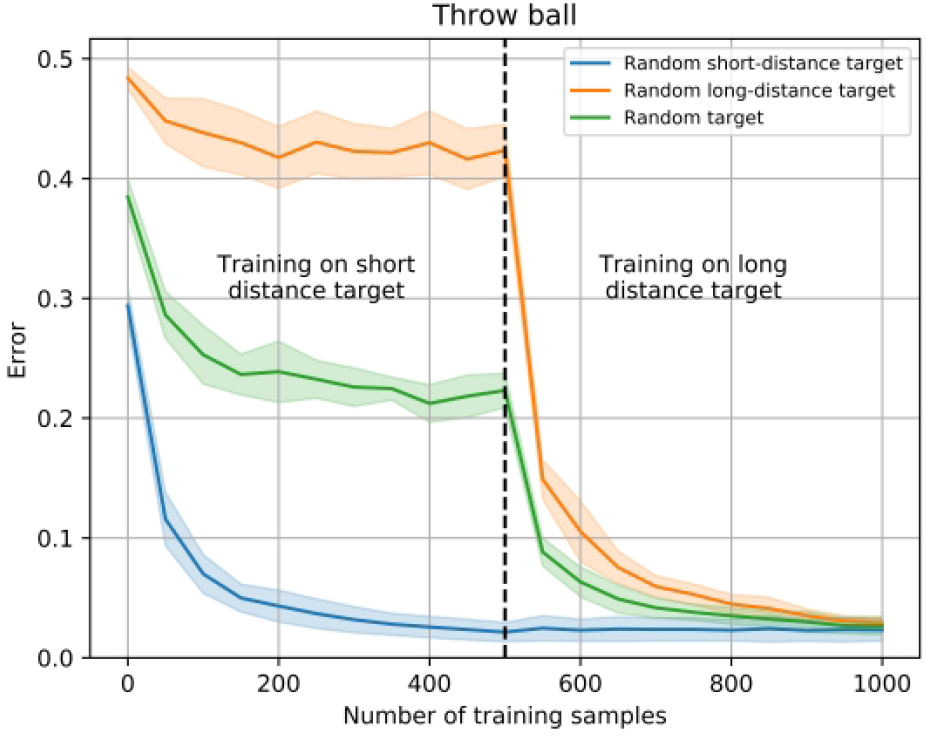
Learning with ordered data samples. We first train the dynamic bump algorithm on 500 samples with short distance targets and then on 500 samples with long distance targets on the throw ball task. We show the performance when only testing on short distance targets (blue line), on uniformly distributed targets (green line) and only on long distance targets (orange line).

In Figure 6, we illustrate the performance of the bump coding scheme on reinforcement learning (RL) tasks. In RL environments an agent should learn to interact with an environment with the goal of maximizing some reward. At any time step, the agent observes the current state of the environment and outputs an action, which in turn affects the state of the environment. The agent obtains rewards for reaching certain states. It is unclear which actions lead to the reward, due to the well known credit-assignment problem. A classical RL method to mitigate this problem is the temporal difference learning method [31], that relies on learning a *policy function* that maps states to actions and a *value function* that maps states to an estimate of the future expected reward. Then, the difference of the estimated expected future reward before and after each action can be computed. Combined with the obtained reward of that time step, one can estimate the reward that arose from that specific action, which allows to update the policy.

**Figure 6:**
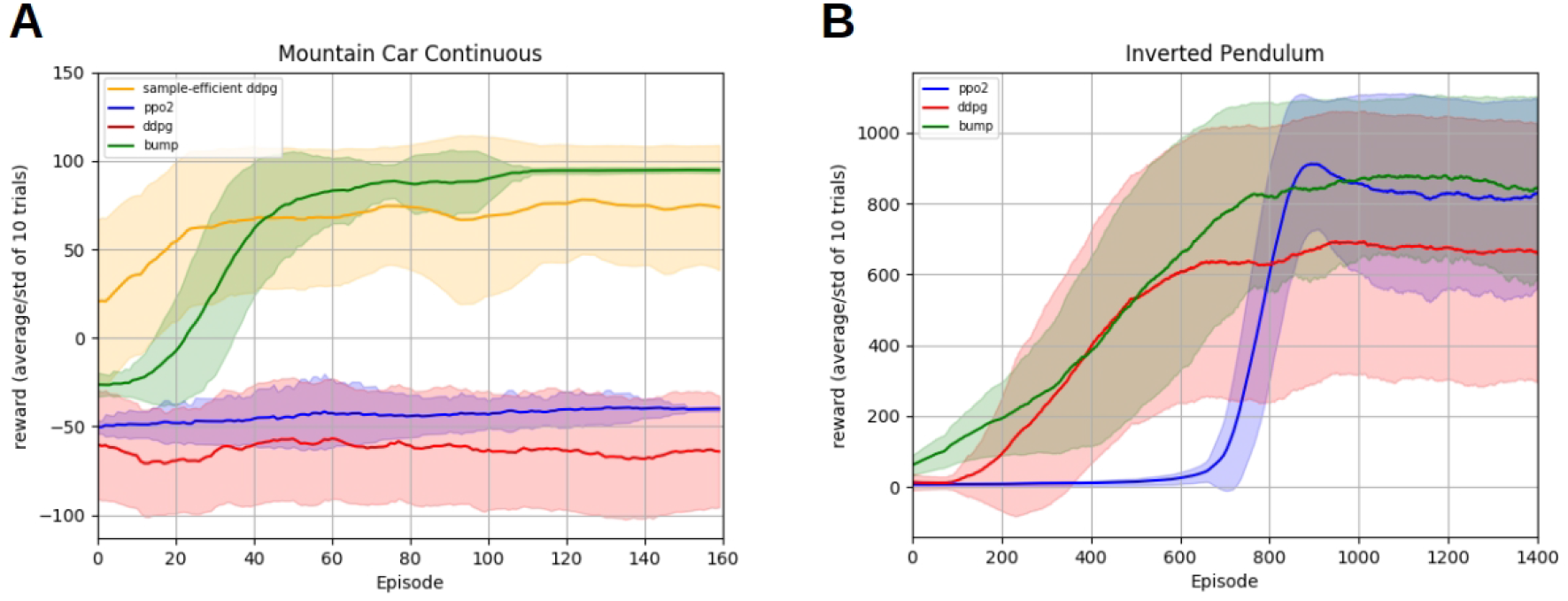
Reinforcement learning experiments. The plots show the learning progress of different learning algorithms on the Mountain Car (A) and Inverted Pendulum (B) task. The tasks are described in more detail in Section 3.5. The RL bump algorithm is compared with the *deep deterministic policy gradient* algorithm (ddpg) and the *proximal policy optimization* algorithm (ppo2), cf. text for implementation details. For every algorithm, the average reward per episode and its standard deviation for 10 different random seeds are smoothed for better readability. The algorithm hyper-parameters are optimized to maximize the mean reward on the last 30 episodes for the Mountain Car, and the last 300 episodes for the Inverted Pendulum, and are given in the supplementary material.

Our *RL bump algorithm* uses the temporal difference learning method. More precisely, it learns the policy with a static bump algorithm, while the value function is stored in tabular representation as done in the literature [31]. At any time step, the temporal difference learning method provides an estimate of the reward arising from the action at this time step. This estimate is used as error feedback for the bump algorithm. Since this estimate might be off, we update the synaptic probabilities more gradually, instead of pruning away synapses immediately as in above algorithms. The magnitude of the updates are chosen proportional to the reward estimates. We refer to Section 3.4 for a detailed description.

We evaluate the RL bump algorithm on two RL tasks, the Mountain Car environment and the Inverted Pendulum, see Section 3.5. The resulting learning curves are displayed in Figure 6. We compare our algorithm with two different deep reinforcement learning algorithms: *deep deterministic policy gradient* (ddpg) [32] and *proximal policy optimization* (ppo2) [33]. Both are state-of-the-art reinforcement learning algorithms that use neural networks as a representation of the policy. For both algorithms, we use the implementation and the standard parameters provided by the OpenAI Baselines [34]. Since these parameters do not aim at efficiently solving the exploration problem of the Mountain Car environment, we added an implementation of ddpg specifically tailored to achieve sample efficiency in the Mountain-Car environment: the parameters and implementation are taken from [35] and the resulting curve is labeled as *sample-efficient ddpg*.

**W**e observe that our algorithm performs comparable or better than the baselines on both environments, see Figure 6. In the Mountain Car experiment, we observe that the OpenAI baselines implementation are unable to make substantial progress on the observed time scale. The sample efficient ddpg implementation from [35] is able to reach higher rewards, but it is very quickly outperformed by our algorithm in terms of final performance. In the Pendulum experiment, we observe a first phase during which the learning curve of our algorithm is very similar to ddpg, whereas ppo2 does not make any progress. In a second phase, the ppo2 learning curve catches up our algorithm, while ddpg is outperformed. We close this section with a word of caution. Figure 5 seems to indicate that our algorithm outperforms deep policy gradient methods for reinforcement learning tasks. However, note that both, Mountain Car and Inverted Pendulum, have a low dimensional input space (2 dimensions for Mountain Car, 4 for Inverted Pendulum). Currently, our algorithm does not scale up to a higher number of dimensions in terms of computational cost, whereas deep policy gradient algorithms have been engineered to deal with high-dimensional spaces.

## 3 Methods

In Section 3.1-3.4, we describe the bump coding scheme and our algorithms formally. Section 3.5, contains a description of all the tasks used for evaluation of the algorithms.

### 3.1 Bump coding scheme

We assume that a population of binary neurons is arranged in a grid. Intuitively, a *d*-dimensional parameter *x* is encoded by a bump of active neurons that lay within the *d*-dimensional cube with side length *k* and center *x*, see Figure 1. We note that our results qualitatively do not depend on the precise shape of the bump, that is, whether it is the *d*-dimensional cube or ball, however our cube shaped bumps facilitate efficient simulations. Formally, a population of *n*^*d*^ neurons encodes values in the *d*-dimensional interval [0, 1]^*d*^. Define *A*_*i*_ = [*n*], where [*n*] denotes the set of integers {1, …, *n*}, and define the set of neurons to be *A* = *A*_1_ × … × *A*_*d*_. Then, a neuron *i* ∈ *A* represents the value (*i*_1_/*n*, … *i*_*d*_/*n*) in [0, 1]^*d*^. Moreover, for *a* ∈ [*n*] define *Int*_*k*_(*a*) to be the set of integers in the interval [*a* − *k*/2, *a* + *k/*2]; to be precise, we actually define *Int*_*k*_(*a*) as the set of integers in the interval [max{0, *a* − *k*/2}, min{*n*, *a* + *k*/2}] in order to take care of cases close to the boundary of the interval. We define the *bump of width k* with center neuron *i* ∈ *A* to be *Int*_*k*_(*i*):= *Int*_*k*_(*i*_1_) × … × *Int*_*k*_(*i*_*d*_). In the bump coding scheme of width *k* the value *a* is encoded by the *Int*_*k*_(*a*)-activation, where an *I*-activation is defined to be the state, where the neurons in *I* are active and the ones not in *I* are inactive.

### 3.2 Network architecture, activation distribution and feedback error measure

In order to learn a mapping 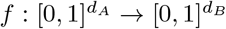, we consider a network consisting of populations *A* and *B* equipped with probabilistic synapses and an abstract continuous attractor mechanism in population *B*. Intuitively, given a bump activation *I* in population *A*, the attractor mechanism activates the bump *J* in population *B* that received most synaptic input from the bump activation in *A*. This way, the probabilistic synapses enable exploration of the output space. More formally, the network consists of population *A* and *B* consisting of 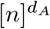 and 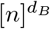 neurons, respectively. These encode values according to the bump coding scheme defined above. They are fully connected by probabilistic synapses with weights *w*_*ij*_ fixed to value 1 and plastic synaptic probabilities *p*_*ij*_. Given an input *x*, the bump 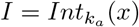 with width *k_A_* and center *x*, the matrix of synaptic probabilites *P* and the bump width *k*_*B*_ in population *B*, we define the following *activation distribution Act*(*I*, *k*_*B*_, *P*) that returns (*J*, *y*), where *J* is the sampled bump in population *B* with center *y*. Assuming that the bump *I* is active in population *A*, we explain now how a bump activation *J* in population *B* is sampled. Denote by 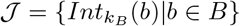 the set of all possible bump activations in *B* with width *k*_*B*_ and by *X*_*ij*_ Bernoulli random variables that are 1 with probability *p*_*ij*_ and 0 otherwise, i.e. they indicate whether synapse (*i*, *j*) is active or not. Then, 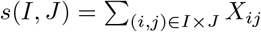 is the total synaptic input that neurons in *J* receive (recall that we assumed that all synaptic weights are constant and equal to 1), and the *abstract continuous attractor mechanism* activates the output bump 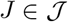 with maximal *s*(*I*, *J*), where ties are broken uniformly at random. Formally, we write (*J*, *y*) ~ *Act*(*I*, *k*, *P*), where 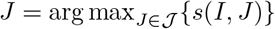 and *y* is the center of bump *J*. In a machine-learning context this can be efficiently implemented by adding a convolutional layer with suitable weights on top of layer *B*. **W**e remark that for the theoretical analysis, we change the activation distribution slightly to be able to deal with the dependencies between distributions *s*(*I*, *J*) and *s*(*I*, *J*′), see supplementary material.

Given a bump *I* in population *A* with center *x* and a sampled activation (*J*, *y*) ~ *Act*(*I*, *k*_*b*_, *P*) the network receives some *error feedback L*(*x*, *y*), where *L* is a function that depends on the output *y* and the target output *f*(*x*), e.g. the euclidean error feedback returns the euclidean distance between output value *y* and the target value *f*(*x*). Note that the learning task is more difficult if the precise definition of the function *L* is unknown to the algorithm. The *error* of a network with learned synaptic probabilities 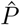 is defined to be the expected error feedback 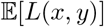, where *x* is chosen uniformly at random in *A* and *y* is sampled according to 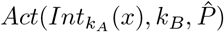.

### 3.3 The static and dynamic bump algorithm

We first explain the algorithms used for the theoretical analysis and then the ones used for the simulations. The goal of these algorithms is to prune away all synapses for every input neuron *x*, except the ones connecting *x* to some small continuous area around *x*’s target neuron *f* (*x*). The basic mechanism to do so is to prune away synapses between the input and output bump whenever the error feedback is larger than some error threshold, as then the target neuron *f* (*x*) is not contained in the output bump.

#### 3.3.1 Algorithms for theoretical analysis

For the following algorithms, every synapse (*i*, *j*) has a synaptic counter *c*_*ij*_ that indicates whether a synapse should be pruned away. Further, the input and output bump widths are set such that the fraction *k*_*B*_/*k*_*A*_ is equal to the Lipschitz constant *C* of the mapping *f* that is to be learned.

The *static bump algorithm (theory)* fixes the input width *k*_*A*_, the output bump width *k*_*B*_ and the error threshold 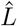 proportional to the desired final error *ℓ* and observes 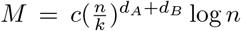 many samples with random input *x*, where 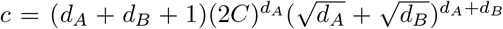. For every sample activation 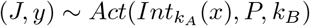, the counters *c*_*ij*_ are incremented by 1 for all synapses between the two bumps 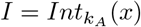 and *J*, if 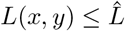 and otherwise decremented by 1. Finally, all synapses with non-positive counters are pruned away. Intuitively, the choice of sample size *M* ensures with high probability^2^ that any input neuron *x* remains to be connected to output neurons *y* with 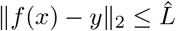.

The *dynamic bump algorithm (theory)* proceeds in phases and repeatedly applies the static version, see Algorithm 4. The bump widths *k* and error threshold 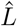 are initially chosen large and divided by 2 in every phase. Intuitively, this causes any input neuron *x* to be connected to a shrinking and shrinking area around target output neuron *f*(*x*).

#### 3.3.2 Algorithms for simulations

The following algorithms can deal with error feedback functions that differ in magnitude for different input regimes and adapt the bump width in a more continuous manner. **W**e give a short overview of the mechanisms used in the algorithms. Firstly, every neuron keeps track of its own error threshold 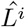. 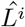 is the running average with decay factor *α* of the error feedbacks obtained when neuron *i* was active. Further, neural counters *c*_*i*_ indicate how accurate the estimates 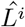 is by counting the number of activations of neuron *i*. Finally, a mechanism to prune long-time inactive synapses is implemented with synaptic counters *d*_*ij*_.

The *dynamic bump algorithm* sets for every sample the input bump width *k*_*A*_ and output bump width *k*_*B*_ both equal to a constant fraction of the number of synapses of neuron *x*. The *static bump algorithm* sets the bump widths to some fixed constant but otherwise proceeds analogously as follows. Any sample activation consists of input and output bumps *I* and *J* with centers *x* and *y*, respectively, and error feedback *L*(*x*, *y*). For each sample, the neurons *i* ∈ *I* update their error threshold according to 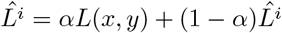 and increase their counter *c*_*i*_ by 1. Synapses (*i*, *j*) between bumps *I* and *J* are pruned away if 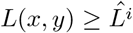 and neuron *i* was active often enough (i.e. *c*_*i*_ ≥ *θ*_*mod*_). Further, synapses (*i, j*) with *i* ∈ *I* and *j* ∉ *J* increase their synaptic counter *d*_*ij*_ by 1 and the ones with *i* ∈ *I* and *j* ∈ *J* reset *d*_*ij*_ = 0. Then, long-time inactive synapses (i.e. *d*_*ij*_ ≥ *θ*_*prune*_) are pruned away. Finally, synapses (*i*, *j*) are consolidated, that is, *p*_*ij*_ is set to 1, if the number of synapses of input neuron *i* drops below a threshold value.

**Algorithm 1:**
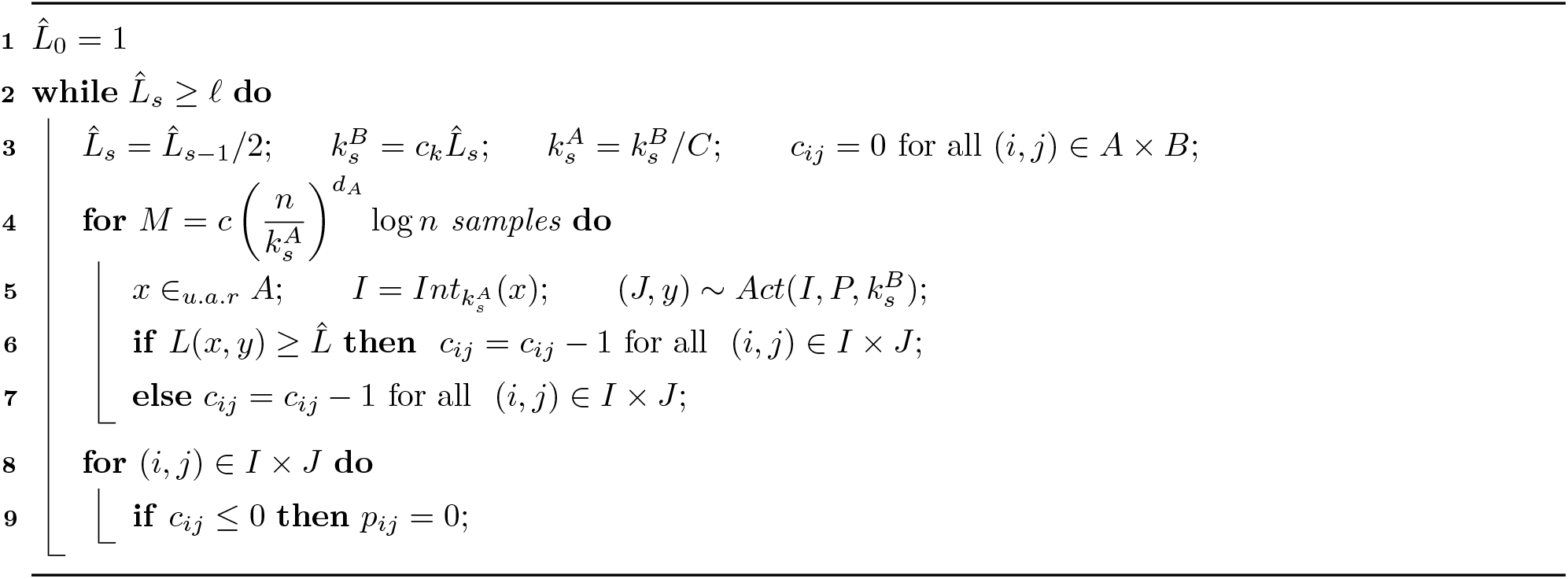
Dynamic bump algorithm for theoretical analysis. It learns a Lipschitz mapping 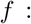 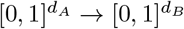. Hyper/parameters: Lipschitz constant or upper bound on Lipschitz constant *C*, desired precision 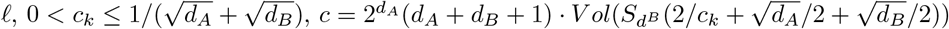, where *Vol*(*S*_*d*_(*r*)) denotes the euclidean volume of the *d*-dimensional ball with radius *r*.

**Algorithm 2:**
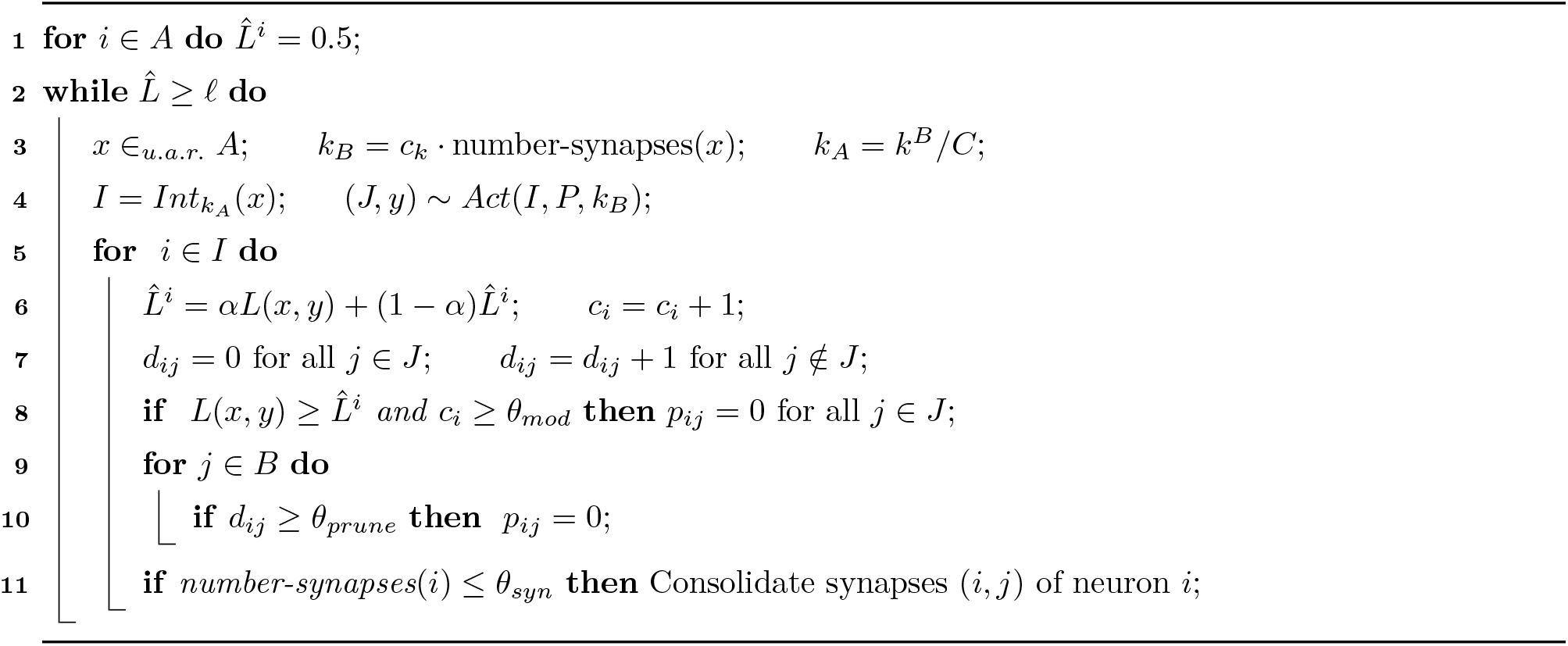
Dynamic bump algorith used for the simulations. It learns a Lipschitz mapping 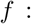 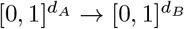. Hyper/parameters: Lipschitz constant or upper bound on Lipschitz constant *C*, desired precision *ℓ*, running average factor *α*, *c*_*k*_, *θ*_*mod*_, *θ*_*prune*_.

### 3.4 Reinforcement learning bump algorithm

In this Section, we describe our *RL bump algorithm*, which combines the classical temporal difference learning method [31] with the bump coding scheme. The building blocks of temporal difference learning are learning a *policy* that maps states to actions and learning a *value function v* that estimates the expected future reward *v*(*S*_*t*_) for states *S*_*t*_. Here, it is assumed that the agent observes the whole state *S*_*t*_ of the environment. To learn these mappings, one computes the *return G*_*t*_, that intuitively is the difference between the estimated future reward *v*(*S*_*t*_) and the real future reward. The *k*-step temporal difference method estimates the real future reward using the bootstrap estimate *v*(*S*_*t*+*k*_) and computes *G*_*t*_ as

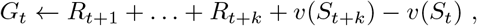

where *R*_*u*_ denotes the reward received at time *u*. Then, the return *G*_*t*_ is used to update the value function, which is stored in a tabular representation [31].

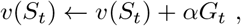

where *α* is a hyper parameter regulating the magnitude of the update. Further, the return *G*_*t*_ is used to update the policy, that in our case is learned with a bump coding algorithm with static bump width that updates the synaptic probabilities gradually. If at time step *t*, the activated bumps were *I* and *J*, we update the probabilities *p*_*ij*_ for all (*i*, *j*) ∈ *I* × *J* according to

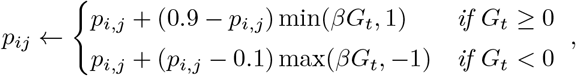

where *β* is a hyper parameter regulating the magnitude of the update. Intuitively, this update increases *p*_*ij*_ proportional to *G*_*t*_ and proportional to the distance of *p*_*ij*_ to 0.9 if *G*_*t*_ ≥ 0, and decreases *p*_*ij*_ proportional to *G*_*t*_ and proportional to the distance of *p*_*ij*_ to 0.1 if *G*_*t*_ ≤ 0. Note that due to the clipping of *βG*_*t*_ to [−1, 1], the invariant is maintained that all probabilities belong to the interval [0.1, 0.9]. The structure of these updates is very similar to policy gradient methods [31], using the reinforce algorithm. However, the reinforce algorithm does not directly apply to our model as it is internally non differentiable, see [17].

### 3.5 Description of tasks

#### 3.5.1 Low dimensional mappings with immediate feedback

As 1-dimensional mappings *f*, we consider the identity function *f*(*x*) = *x*, a sinus function *f*(*x*) = sin(*x*) and a second order polynomial *f* (*x*) = *x*^2^ − 3*x* + 1. For all functions, we consider the absolute distance *L*(*x*, *y*) = |*f*(*x*) − *y*| as error feedback, where *x* and *y* are input and output of the network.

In the *throw ball task*, the network has to learn to throw a ball to a certain distance. The target distance is given as a 1-dimensional input to the network. The networks gives a 2-dimensional output, that consists of the vertical throwing angle and the initial speed of the ball. The error feedback is the absolute difference between the distance where the ball touches the ground and the target distance. Note that the optimal output is underdetermined, as for any angle in (0, π/2) there exists a speed such that any target distance can be hit.

In the *robotic arm task*, the network learns how to control a simple robotic arm with two degrees of freedom. The arm is composed of two rigid moving parts and is connected to a fixed anchorage point. As we restrict to movements in the plane, the agent only has to control angles, one at each joint. The target position is given as input in Cartesian coordinates and the network outputs two angles that are applied to the two joints. The feedback is given by the euclidean distance between the actual position of the arm and the target position.

#### 3.5.2 Reinforcement learning tasks

The RL environments used for assessing the RL bump algorithm are from the OpenAI Gym [36] toolkit: the continuous version of the classical Mountain Car control problem (MountainCarCountinuous-v0), and the Inverted Pendulum environment from the MuJoCo suite (InvertedPendulum-v2). In the *mountain car task* the goal is to reach the top of a hill, that can only be reached by obtaining momentum when driving down the neighbouring hill. The network receives as input the position and speed of the car and outputs the acceleration that is applied to the car. There is a large positive reward if the car reaches the top of the hill and a small negative reward for the fuel use in every time step. In the *inverted pendulum task* the goal is to balance an inverted pendulum on a cart. The network receives 4-dimensional input describing position and velocity of the cart and pendulum, and it outputs the acceleration applied to the cart. As long as the pendulum does not fall to the ground, there is a positive reward in every time step.

## 4 Discussion

In this section, we first relate our work to related work from the field of machine learning that studies sample efficient learning, then we discuss the bio-plausibility of our proposed coding scheme and learning algorithms, and finally we discuss the insights gained about the bio-plausible mechanisms used in this study.

### 4.1 Sample efficient learning in machine learning

How to improve sample efficiency of learning algorithms is a major topic in the field of machine learning in general and the field of reinforcement learning in particular [37]. Data samples are often limited, and training of artificial networks is computationally costly. A common approach to improve sample efficency is to handcraft artificial networks to the task at hand. The most famous example are convolutional neural networks [38, 39], where the translational invariance property of images is hand-wired into the convolutional network architecture. Another successfull approach is to store all observed samples or the neural states that encode these samples. Then, inputs are classified according to the most similar samples in storage. This idea is present in non-parametric approaches such as the nearest neighbour methods [40], as well as in deep neural networks augmented with external memory systems [41] and attention mechanisms [42, 43]. In reinforcement learning this idea is known as episodic reinforcement learning [44, 45, 46, 47]. Further, an approach to improve sample efficiency is meta learning [48], which is also often referred to as ‘learning to learn’. In the meta learning setting, an outer learning system adjusts parameters or learning mechanisms of an inner learning system in order to improve the performance and efficiency of the later [49, 50, 51, 52, 53]. The outer learning system usually performs updates in a slow timescale, whereas the inner learning system can adapt fast to new environments, e.g. evolutionary algorithms can optimize learning architectures or loss functions to improve their sample efficiency [54, 55].

Our approach is orthogonal to all these approaches. In essence, our work shows that the coding scheme of a network affects its sample efficiency and that adapting the coding scheme during learning can improve its sample efficiency.

### 4.2 Bio-plausibility of our coding scheme and learning mechanisms

The proposed coding scheme and learning algorithms are of abstract nature and we do not intend to argue that they might be implemented in biological systems precisely in this form. However, we do claim that neural implementations of the basic concepts used by our model are plausible and the brain might use similar mechanisms for computation.

Our primary assumption that information is encoded and processed by populations of tuned neurons is supported by the abundance of such neurons across brain areas [3, 4, 5, 6, 7]. The bump coding scheme described in Section 3 requires that geometrically close-by neurons have close-by preferred stimuli parameters. Such geometrically ordered networks are indeed present in real neural network, such as the drosophila fly compass system [8, 9]. The underlying network-wiring that gives rise to bump-like activation patterns is generally believed to follow the circuit-motiv of local excitation and long-range inhibition, as suggested by experimental evidence [9] and theoretical findings [10, 11, 12]. Note however, that the geometrical ordering of the neurons is not necessary for the results presented in this work. Indeed, the geometrical arrangement can be arbitrary if network-wiring supports activity patterns consisting of neurons with similar preferred stimuli parameters. Such network wiring consists of excitatory connections between neurons that are active for similar stimuli and inhibition that limits the total activity. It can be found across brain areas and animal species [56, 57, 58], and is often assumed by theoretical studies [12, 13, 59]. It is conceivable that such experimentally observed wiring motives implement a version of the abstract continuous attractor mechanism used in this paper.

The dynamic bump algorithm requires a dynamic adaptation of the bump width during the learning process. Experimental and theoretical studies give evidence that the tuning curve width is controlled by inhibition [60, 61, 59]. Thus, controlling the strength of inhibition in the system yields a straight forward explanation of how the bump width could be adjusted during the learning progress.

Moreover, the assumption of constant weights and binary neurons are mere abstractions for mathematical simplicity. Due to the on-off nature of binary neurons, we approximated the bell shaped tuning curves by rectangular tuning curves. It seems plausible that the results would translate qualitatively to networks of rate neurons with bell shaped tuning curves. Stable bump like activity patterns also can be produced by spiking networks [14]. We leave extensions of our algorithms that are more bio-plausible for future investigations. Moreover, we note that all results from this work also hold if the populations are sparsely instead of fully connected. As long as the bump width is broad enough, sufficiently large population codes give rise to stable learning mechanisms for sparsely connected populations [62].

Furthermore, our plasticity rules are plausible in the sense that they solely depend on pre- and post-synaptic activity, a global reward feedback and memory traces of these quantities. All the neuronal and synaptic counters used in our algorithms require only local storage of activity and reward feedback traces.

### 4.3 Functional role of tuning curve width

A large body of literature in theoretical and experimental neuroscience investigated tuning curve shape under the aspect of optimal coding [63, 64, 65, 66, 67, 68, 69, 22, 70, 71, 72, 73, 74, 75, 76, 77, 78, 79, 80]. Also the width of tuning curves was analysed from an information theoretical viewpoint. [81] and [82] established a dependence between optimal tuning width and the dimensionality of the encoded parameter, [83] and [84] showed that optimality of tuning width heavily depends on the level of noise and covariance of the noise in the system, and [85] found that optimal tuning width depends on the prior uncertainty and on the length of the decoding time window. Such studies can explain the sharpening of tuning curves that is observed in a variety of experimental set-ups [86, 87, 88, 89, 90].

In this work, we introduce sample efficiency as a novel notion of optimality. If the neural code is optimized for sample efficient learning, then the model analyzed in this paper predicts that the tuning curves sharpen during the process of learning. In fact, this phenomena is known to occur in the inferiour temporal cortex, where tuning curves of shape selective neurons sharpen during acquaintance to new objects [91, 92], as well as in many sensory areas during development [93, 94, 95]. The precise relation between tuning curve width and sample efficiency likely depends on the applied plasticity mechanisms. Nonetheless, for any plasticity mechanism that requires pre- and post-synaptic activity, the tuning curve width yields an upper bound on the number of synaptic weight updates, because it limits the number of active neurons per sample. Therefore, the basic principle that larger tuning curve width leads to more synaptic updates per sample and thus faster learning, may apply to many plasticity mechanisms.

### 4.4 Functional role of probabilistic synapses

The functional role of probabilistic synapses is highly debated [15]. The proposed functional roles include regularization and improved generalization in deep neural networks [96, 97] and energy saving constraints [98]. Further, probabilistic synapses can give rise to a good exploration explotation trade-off in reinforcement learning [97, 99, 100], and synaptic sampling can be seen as sampling from some posterior distribution [101, 102, 100]. Our model is in line with the last two proposals. In our model, probabilistic synapses combined with an continuous attractor mechanism encode the uncertainty of the learned input-output mapping and implement the variability and exploration that is required for reward-based learning.

### 4.5 Conclusion

In this work, we asked how sample efficient learning is affected by the neural coding scheme. We showed that population codes with tuned neurons support sample efficient learning for low dimensional tasks with immediate reward feedback and low dimensional reinforcement learning tasks. For these tasks, our gradient-free learning algorithm is competitive to multi-layer perceptrons trained by backpropagation. These findings might inspire an integration of tuning curve coding schemes into machine learning approaches, especially, if data-samples are limited and no access to gradient information is given. For our learning mechanisms, we found that tuning curve width severely influences the sample efficiency. We showed that for static tuning widths, there is a trade-off between sample efficiency and final precision. Broad tuning curves give rise to sample efficient learning, whereas narrow tuning curves account for high final precision. Moreover, we showed that dynamic adaptation of the tuning width results in both high sample efficiency and high final accuracy. These results propose sample efficient learning as a functional role of the tuning curve width.

## Author Contributions

FM, AS: development of model set-up and research question, design, development and analysis of learning algorithms, writing of the manuscript. RD: implementation and simulation of the learning algorithms, development of the RL-bump algorithm, proofreading of the manuscript.

## Acknowledgments

We thank Ulysse Schaller for preliminary work on the model set-up and research question during his master thesis.

## Data Availability Statement

The code used for the simulations of the learning algorithms is provided under the following link [https://github.com/rdang-nhu/Sample_Efficient_Tuning_Curves].

## Supplementary Material

### A Mathematical analysis

#### A.1 Formal statement of the Theorems

We first restate the theorems from the results section of the main paper in a slightly more formal way. For simplifying the notation in the proofs, we rescale the domain and codomain of the function *f*. Let 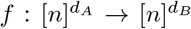, where [*n*] denotes the set of integers {1, …, *n*}. This rescaling from [0, 1] to [1, *n*] is convenient as in this way neuron *i* ∈ [*n*] represents value *i*. Note that this also rescales the error. An error smaller than *ℓ* implies an error smaller than *ℓ/n* in the 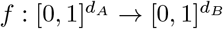 case. For completeness we also state the static bump algorithm (theory) formally in Algorithm 3 and restate the dynamic bump algorithm (theory) in Algorithm 4.

**Algorithm 3:**
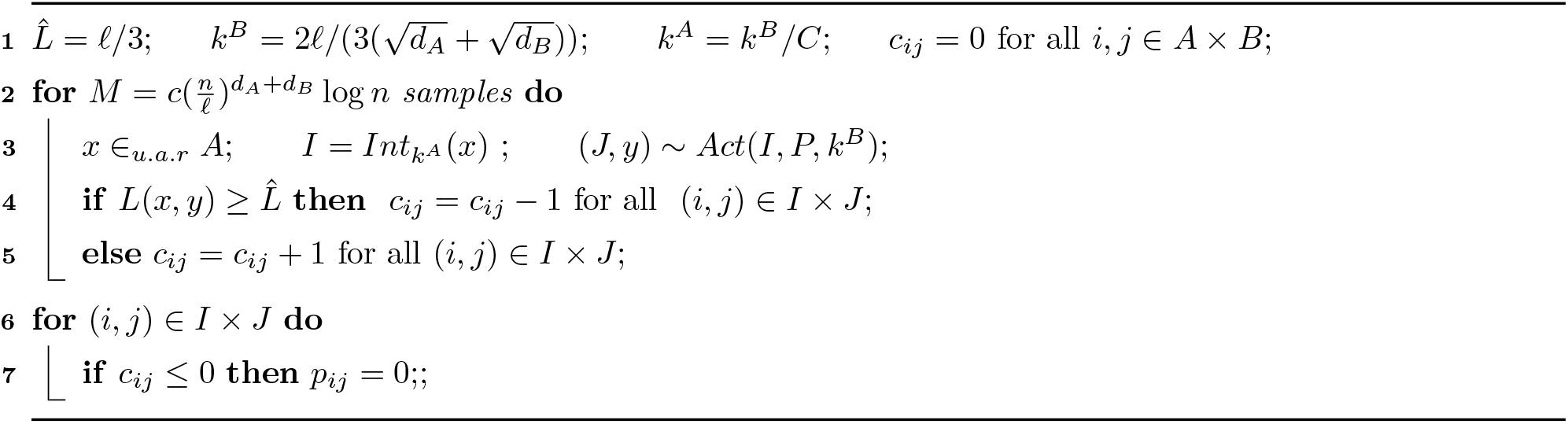
The static bump algorithm (theory) for learning a Lipschitz function 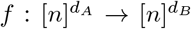. Hyper-parameters: Lipschitz constant or upper bound on Lipschitz constant *C*, desired precision *ℓ*, 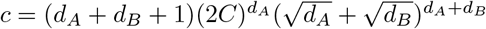.

**Algorithm 4:**
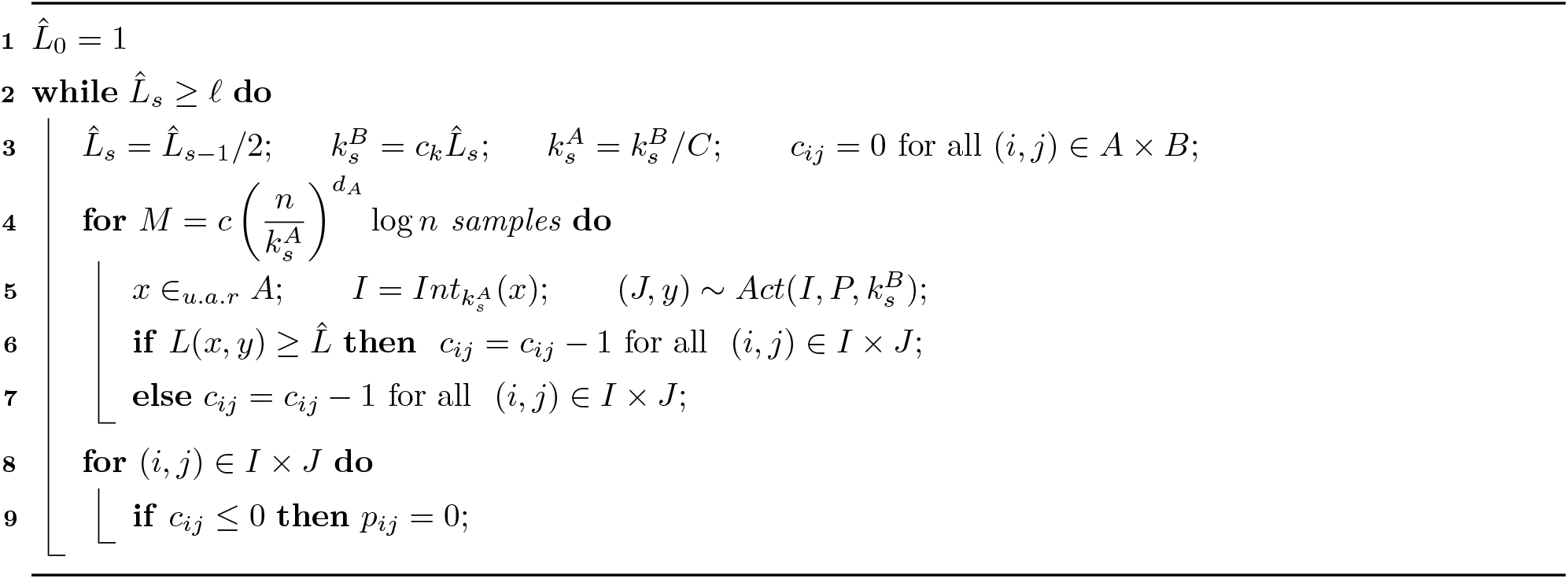
The dynamic bump algorithm (theory) for learning a Lipschitz function 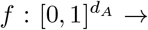 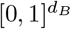. Hyper-parameters: Lipschitz constant or upper bound on Lipschitz constant *C*, desired precision 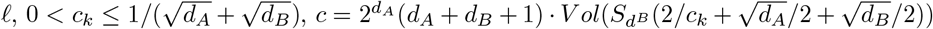, where *Vol*(*S*_*d*_(*r*)) denotes the euclidean volume of the *d*-dimensional ball with radius *r*.

##### Theorem 4 (Static bump width *k*).

*Let* 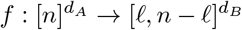 *be a C-Lipschitz continuous function, where ℓ is the desired precision. Assume that the error feedback L*(*x*, *y*) = ‖*f*(*x*) − *y*‖_2_ *is the Euclidean distance between output y and target output f*(*x*). *Then*, *Algorithm 3 achieves with high probability error smaller than ℓ after* 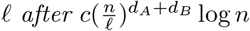 *samples, where c is as stated in Algorithm 3*.

We remark that the restriction, that *f* maps not too close to the boundary of 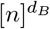 is required since the neurons close to the boundary have less neighbouring neurons and thus it is rather unlikely that their surrounding interval receives most synaptic input.

##### Theorem 5 (Dynamic bump width *k*).

*Let c*_*k*_ > 0. *Let* 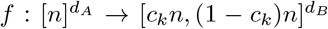 *be a C-Lipschitz continuous function. Assume that the error feedback L*(*x*, *y*) = ‖*f*(*x*) − *y*‖_2_ *is the Euclidean distance between output y and target output f*(*x*). *Then*, *Algorithm 4 achieves with high probability errors smaller than ℓ after* 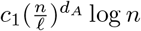 *samples, where* 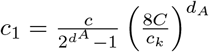 *and c and c*_*k*_ *are defined in Algorithm 4*.

We also restate the lower bound on the sample efficiency:

##### Theorem 6 (Lower Bound).

*Let C* > 0. *There exist C-Lipschitz functions* 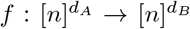, *such that any Algorithm A, that receives for every sample only a constant number of bits as feedback, requires at least* 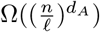 *samples*^3^ *to achieve accuracy ℓ*.

#### A.2 Notation and preliminary lemmas

In the paper, we defined the total synaptic input from bump *I* to bump *J* as 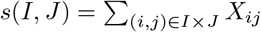, where the *X*_*ij*_ are Bernoulli random variables that are 1 with probability *p*_*ij*_. Intuitively, the distributions *s*(*I*, *J*) and *s*(*I*, *J*′) are identical if *I* has identical synaptic probabilities to *J* and *J*′, and therefore, the activation probabilities of bump *J* and bump *J*′ should be the same. But if *J* and *J*′ overlap, then *s*(*I*, *J*) and *s*(*I*, *J*′) are not independent which would complicate the theoretical analysis. To overcome this issue, we slightly change the activation procedure here. We define *s*(*I*, *J*) to be the expected total synaptic input 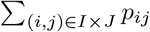. Then, given an activation *I* in population *A*, we sample the output activation *J* according to an adapted softmax probability measure 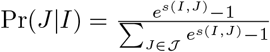. Note that this probability measure is well defined as *I* has at least one outgoing synaptic probability larger than 0. In addition, it has the properties that if all synapses in *I* × *J* are zero, then Pr(*J*|*I*) = 0, and if *s*(*I*, *J*_1_) = *s*(*I*, *J*_2_) then Pr(*J*_1_|*I*) = Pr(*J*_2_|*I*).

We now define some notations that are convenient for the analysis.

- Recall that for *x* ∈ [*n*]^*d*^, we defined *Int*_*k*_(*x*) = *Int*_*k*_(*x*_1_)× … ×*Int*_*k*_(*x*_*d*_), where *Int*_*k*_(*x*_*i*_) = [max{0, *x*_*i*_− *k*/2}, min{*n*, *x*_*i*_ + *k*/2}].
- An (*I*, *J*, *x*, *y*)-activation consists of bumps *I* and *J* with centers *x* and *y*, respectively.
- For an (*I*, *J*, *x*, *y*)-activation, we define *R*_*k*_(*x*, *y*) to be the set of synapses between bump *I* and bump *J*. Formally, we define 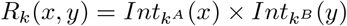, where *k* denotes the tuple (*k*^*A*^, *k*^*B*^). Note that synapse (*i*′, *j*′) being in *R*_*k*_(*i*, *j*) is equivalent to synapse (*i*, *j*) being in *R*_*k*_(*i*′, *j*′).
- We call *R*_*k*_(*i*, *j*) *full* if it contains 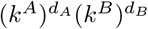 many synapses (*i*, *j*) with *p*_*ij*_ = 1/2 and *empty* if it contains no synapses with *p*_*ij*_ > 0. Otherwise, we call *R*_*k*_(*i*, *j*) *moderate*.
- Further, for a given input neuron *i*, we define 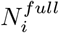, 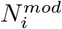 and 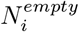 as the number of output neurons *j* for which *R*_*k*_(*i*, *j*) is full, moderate and empty, respectively.
- Define *p*(*j*|*i*) = Pr(*J*|*I*) to be the probability that *j* is the center of the output bump *J* conditioned on *i* being the center neuron of the input bump *I*.

Our first lemma states some bounds on the values *p*(*j*|*i*) that will be useful for the proof.

##### Lemma 7 (Bounds on *p*(*j*|*i*)).

*The following bounds on the probabilities p(j|i) hold*.

1. *If R*_*k*_(*i*, *j*) *is empty, then p*(*j*|*i*) = 0.
2. *Consider a fixed i* ∈ *A*. *For all j with full R*_*k*_(*i*, *j*), *p*(*j*|*i*) *has the same value* 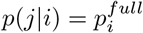. *It holds that* 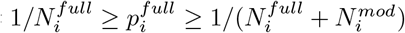.
3. *If R*_*k*_(*i*, *j*) *is moderate, then* 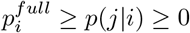.

*Proof*. Let 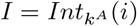 and 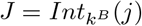. Recall that 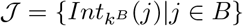 is the set of possible output activations, and that 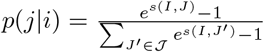. Note that 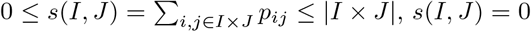 holds if and only if 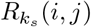 is empty, and *s*(*I*,*J*) = |*I* × *J*| holds if and only if 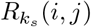 is full. From this, the statements of the lemma follow easily.

#### A.3 Static bump width *k*

Let us first outline the proof of Theorem 4, before giving the details of proof later in this section. We call a synapse good, if it is inside and not too close to the boundary of the red area in Figure 1 and bad if it is outside and not to close to the boundary of the red area. Formally, a synapse (*i*, *j*) is *good* if 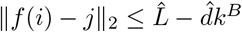 and *bad* if 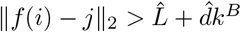, where 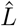 is the error feedback threshold and 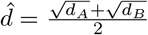. We show that good synapses (*i*, *j*) can only receive positive counter updates. The synapse receives at least one counter update if *j* is not to close to the boundary of *B*, which implies that *p*_*ij*_ = 1/2 after execution of the algorithm. For bad synapses (*i*, *j*), we show that they only receive negative counter updates, which implies that *p*_*ij*_ = 0 after execution of the algorithm. This ensures that the error is at most *ℓ*. Formally, we show that the following 5 properties hold.

1. Good synapses (*i*, *j*), i.e. 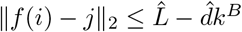, obtain only positive counter updates.
2. Bad synapses (*i*, *j*), i.e. 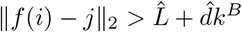, obtain only negative counter updates.
3. 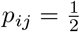 for all good synapses with 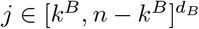 after execution of the algorithm.
4. *p*_*ij*_ = 0 for all bad synapses after execution of the algorithm.
5. The error is at most *ℓ* after execution of the algorithm.

##### Property 1

Let (*i*, *j*) be such that 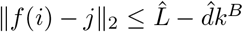, and let (*I*, *J*, *i*′, *j*′) be an activation such that synapse (*i*, *j*) is active, that is, (*i*, *j*) ∈ *R*_*k*_(*i*′, *j*′). Note that the diagonal of bumps *I* and *J* have length 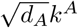 and 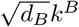, respectively. This implies that 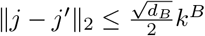, and the Lipschitz continuouity of *f* implies that 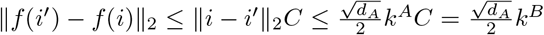, where the last equality follows from the choice of *k*^*A*^ and *k*^*B*^ in Algorithm 3. The triangle inequality implies

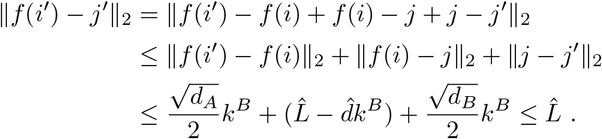

Therefore, synapses (*i*, *j*) is only updated for samples with error feedback 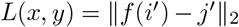 smaller than 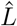, which implies that the synapse only receives positive counter updates.

##### Property 2

This statement follows similarly as the last one. Let (*i*, *j*) such that 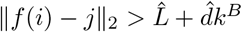, and let (*I*, *J*, *i*′, *j*′) be an activation such that synapse (*i*, *j*) is active. Then, (*i*, *j*) ∈ *R*_*k*_(*i*′, *j*′) implies that 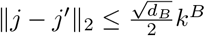 and 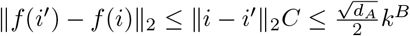. The triangle inequality implies

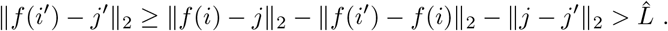

It follows that synapse (*i*, *j*) receives only negative counter updates.

##### Property 3

Let (*i*, *j*) be a good synapse with 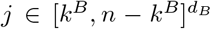. Property 1 states that *c*_*ij*_ always increases when synapse (*i, j*) is active. Thus, *c*_*ij*_ ≥ 1 holds if the synapse is active at least once during the *M* samples. In the following, we show that this happens with high probability. Note that during the execution of the algorithm we have *p*_*ij*_ = 1/2 for all (*i*, *j*) ∈ *A* × *B*. Therefore, *R*_*k*_(*i*, *j*) is full for all 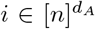 and 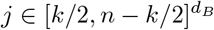, and Lemma 7 implies that *p*(*j*|*i*) is maximal and equal to 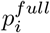 for these values of *i* and *j*. Denote by 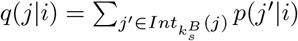 the probability that output neuron *j* is active given that *i* is the center neuron of the input. Then 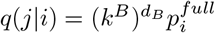 for 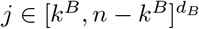. Note that the probability that *i*′ is the center neuron of the input is 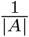. Thus, the probability that synapse (*i*, *j*) is active in a fixed round is

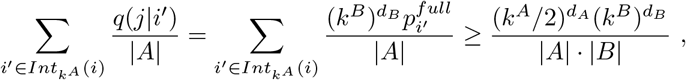

where the inequality follows from 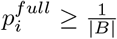 and 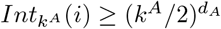 (this lower bound comes from the case that *i* is in the corner of the input space *A*). Using the inequality 1 − *x* ≤ *e*^−*x*^, it follows that the probability that synapse (*i*, *j*) does never turn active during the 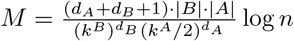 rounds is smaller than

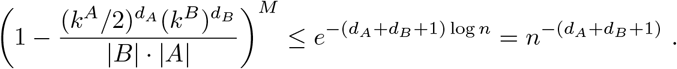

Then, a union bound over all 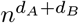 synapses implies that Property holds 3 with probability 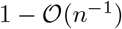.

##### Property 4

Property 2 sates that *c*_*ij*_ cannot be increased for bad synapses. Therefore, *c*_*ij*_ ≤ 0 and *p*_*ij*_ = 0 hold after execution of the algorithm.

##### Property 5

By Property 4 it follows that *p*_*ij*_ = 0 for 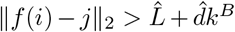. The triangle inequality implies that *R*_*k*_(*i*, *j*) is empty for 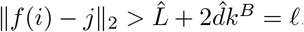, where the last equality follows from the choice of 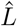 and *k*^*B*^ in Algorithm 3. Thus, by Lemma 7, *p*(*j*|*i*) = 0 if 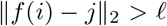, which implies that the error is smaller than *ℓ*.

##### A.4 Dynamic bump width *k*

In this section we proof Theorem 5. The proof ideas are analogous to the proof of Theorem 4 in the last section. We will show that similar properties as in the last section hold after each phase of Algorithm 4. In phase *s*, we call a synapse (*i*, *j*) *good* if 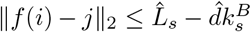, and we call it *bad* if 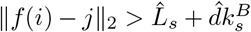 where 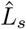 is the error feedback threshold of phase *s* and 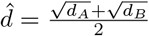. We show that good synapses (*i*, *j*) only receive positive counter updates and that *p*_*ij*_ = 1/2 holds after phase *s* if *j* is not too close to the boundary of *B*. Note that this boundary condition is more complicated than for the static bump algorithm, because of the iterative application of the static algorithm, c.f. Property 3 below. Bad synapses (*i*, *j*) only receive negative counter updates, which implies that *p*_*ij*_ = 0 after phase *s* of the algorithm. This implies that the error is at most 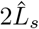 after phase *s* of the algorithm. Formally, we show that the following 7 properties hold. We remark that the purpose of Properties 5 and 6 is to facilitate the analysis of the next phase.

1. Good synapses, i.e. 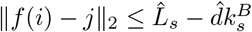, receive only positive counter updates during phase *s*.
2. Bad synapses, i.e. 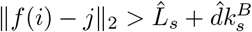, receive only negative counter updates during phase *s*.
3. After execution of phase *s*, 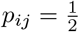 holds for all good synapses (*i*, *j*) with 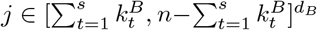.
4. After execution of phase *s*, *p*_*ij*_ = 0 holds for all bad synapses.
5. If 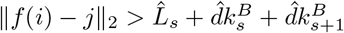, then 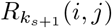 is empty after execution of phase *s*.
6. Let *α* > 0. If 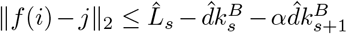 and 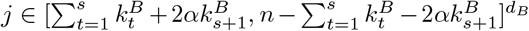 then 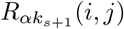 is full after execution of phase *s*
7. The error is at most 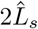 after execution of phase *s*.

Let us now show how these properties imply the Theorem. Recall that 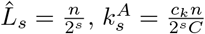 phase *s* requires 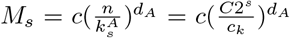 many samples. Property 7 implies that the algorithm reaches error smaller than *ℓ* in the phase 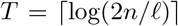 for which 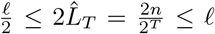 holds, which is equivalent to 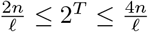. Therefore, the total number of samples required by the algorithm is

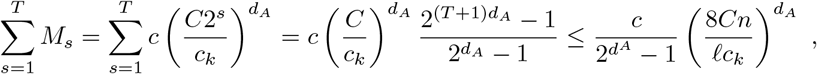

where we used the formula for a sum of a geometric series. This implies Theorem 5.

It remains to prove Properties 1-7. The proof follows by induction over *s*. Observe that before phase 1 (i.e. *s* = 0, *L*_0_ = *n*, 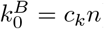) all properties are satisfied. Given that all properties are satisfied after phase *s* − 1, we show in the induction step that all properties are satisfied with probability 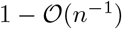 after phase *s*. It follows that after phase 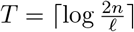 all properties are satisfied with probability 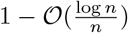.

###### Property 1

This statement follows analogously to Property 1 in Section A.3 by the Lipschitz continuity of *f* and triangle inequality.

###### Property 2

The proof is analogous to the proof of Property 2 in Section A.3.

###### Property 3

Similar to the proof of Property 3 in the last section, we prove that that *c*_*ij*_ ≥ 1 holds with probability 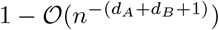 for such synapses. Then, by union bound over all synapses satisfying the assumptions of 3, it holds with probability 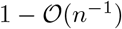 that *c*_*ij*_ ≥ 1 for all these synapses.

Let (*i*, *j*) be a synapse satisfying the assumptions of Property 3

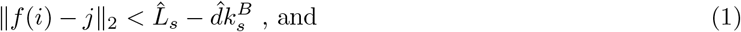

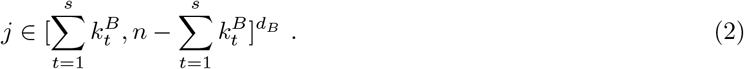

The choice of *c*_*k*_ in the algorithm implies that that 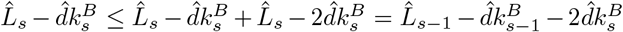.Then, (2) and Property 6 for phase *s* − 1 with *α* = 2 imply that 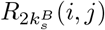 is full. Therefore, 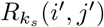 is full for 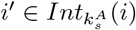 and 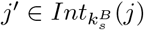, and Lemma 7 implies that 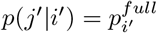.

By Property 1, good synapses can only receive positive counter updates. Thus, *c*_*ij*_ ≥ 1 holds if (*i*, *j*) is active at least once during the *M* rounds. Denote by 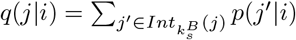 the probability that output neuron *j* is active given that *i* is the center neuron of the input. Note that the probability that *i*′ is the center neuron of the input is 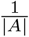. Thus, the probability that synapse (*i*, *j*) is active for a fixed sample is

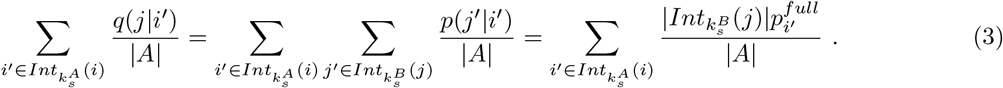

Property 5 from phase *s* − 1 implies that if 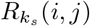 is full or moderate, then *j* lies in the ball with center *f* (*i*) and radius 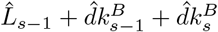. By Lemma 7, it follows

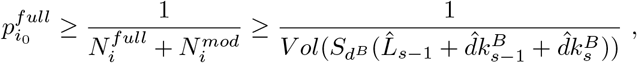

where 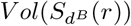 denotes the euclidean volume of the *d_B_* dimensional ball with radius *r*. Note that 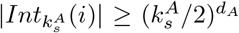 (this lower bound comes from the case where *i* is in the corner of input space *A*), 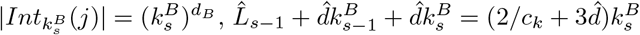 and 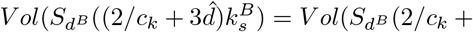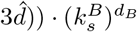. Thus, the probability from Equation (3) that synapse (*i*, *j*) is active for a fixed sample is at least

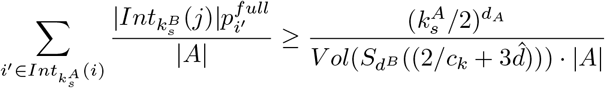

By the inequality 1 − *x* ≤ *e*^−*x*^, it follows that the probability that synapse (*i*, *j*) does never turn active during the 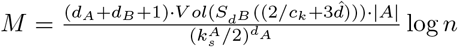 rounds is smaller than

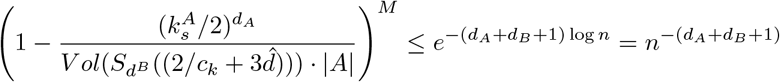

which proves the claim.

###### Property 4

For bad synapses (*i*, *j*), Property 2 states that *c*_*ij*_ can only be decreased in phase *s*. This implies that *c*_*ij*_ ≤ 0 and *p*_*ij*_ = 0 after phase *s*.

###### Property 5

For 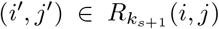 the Lipschitz continuity of *f*, the assumption 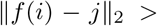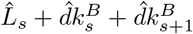 and the triangle inequality implies

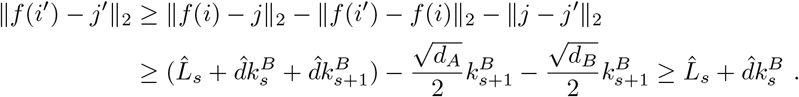

The statement follows by Property 4.

###### Property 6

Let *α* > 0 and (*i*, *j*) such that the assumptions are satisfied. Let 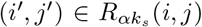. The Lipschitz continuity and the triangle inequality imply that synapse (*i*′, *j*′) is good:

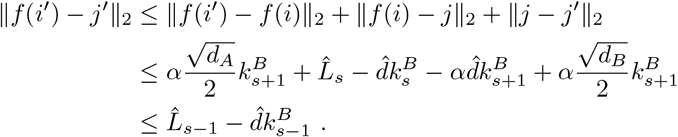

Moreover, 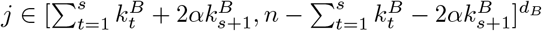 and 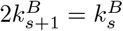 imply that 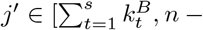 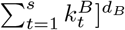. By Property 3, it follows that 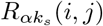 is full.

###### Property 7

By Property 5 and Lemma 7, it follows that 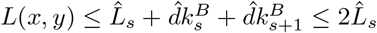, where the last inequality follows from the choice of 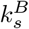 in the algorithm.

**Table 1:** Algorithm Hyper-Parameters for the Reinforcement Learning Experiments

| Parameters | MountainCarContinuous v0 | InvertedPendulum v2 |
| --- | --- | --- |
| Input grid dimensions | 50x50 | 20x20x20x20 |
| Output grid dimensions | 100 | 50 |
| Policy learning rate | 0.1 | 0.003 |
| Value function learning rate | 0.5 | 0.5 |
| Initial value function | 1 | 140 |
| K input | 5 | 3 |
| K output | 5 | 7 |

##### A.5 Lower bound

In this section we prove Theorem 6. Let *d*_*A*_ be the input dimension. Let us construct a Lipschitz function 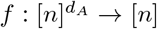 with Lipschitz constant *C* > 0 for which 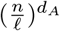 bits of information are required in order to approximate it with precision *ℓ*. Let *m* = ⌈2*ℓ*/*C*⌉. Consider the following family 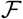 of Lipschitz functions. For simplicity assume *n* is divisible by *m*. For 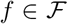 it holds that *f*(*x*) = 0 if *x*_*i*_ ≡ 0 mod 2*m* for some *i*, and *f*(*x*) ∈ {−2*ℓ*,2*ℓ*} if *x*_*i*_ ∈ {*m*, 3*m*, 5*m*, …} for all *i*. For the points *x* for which *f*(*x*) is not defined yet, we take a linear interpolation between the points defined above. This way the slope of any linear part is at most 2*ℓ*/*m* ≤ *C*. Now, let us draw a 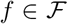 uniformly at random. In order to achieve precision *ℓ* on the function *f*, one needs to know for each point on the grid 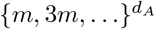 whether the function value is 2*ℓ* or −2*ℓ*. Therefore, 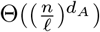 bits of information are required. Since the algorithm can obtain at most a constant number of bits of information per sample, 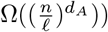 samples are required. If the codomain of the function is multi-dimensional, this lower bound holds as well, by only considering the first output dimension.

### B Details and hyperparameters for experiments

Table 1 shows the hyper parameters used to obtain the results of the RL bump algorithm illustrated in Figure 5 of the main paper.

## Footnotes

1 A function *f* is Lipschitz continuous with Lipschitz constant *C*, if |*f* (*x*) − *f* (*y*)| ≤ *C*|*x* − *y*| for all *x, y*. Intuitively, this is the case if the slope of *f* is everywhere smaller than *C*.

2 *With high probability* means with probability tending to 1 as *n* tends to ∞.

3 We use Landau notation 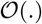, Θ(.), Ω(.),… to denote the asymptotic behaviour with respect to *n* → ∞

